# ChEC-seq kinetics discriminates transcription factor binding sites by DNA sequence and shape *in vivo*

**DOI:** 10.1101/027334

**Authors:** Gabriel E. Zentner, Sivakanthan Kasinathan, Beibei Xin, Remo Rohs, Steven Henikoff

**Affiliations:** Basic Sciences Division, Fred Hutchinson Cancer Research Center, Seattle, WA 98109, USA; Medical Scientist Training Program, University of Washington School of Medicine, Seattle, WA 98195, USA; Molecular & Cellular Biology Graduate Program, University of Washington School of Medicine, Seattle, WA 98195, USA; Molecular and Computational Biology Program, Departments of Biological Sciences, Chemistry, Physics, and Computer Science, University of Southern California, Los Angeles, CA 90089, USA; Howard Hughes Medical Institute, Fred Hutchinson Cancer Research Center, Seattle, WA 98109, USA

## Abstract

Chromatin endogenous cleavage (ChEC) uses fusion of a protein of interest to micrococcal nuclease (MNase) to target calcium-dependent cleavage to specific genomic loci *in vivo.* Here we report the combination of ChEC with high-throughput sequencing (ChEC-seq) to map budding yeast transcription factor (TF) binding. Temporal analysis of ChEC-seq data reveals two classes of sites for TFs, one displaying rapid cleavage at sites with robust consensus motifs and the second showing slow cleavage at largely unique sites with low-scoring motifs. Sites with high-scoring motifs also display asymmetric cleavage, indicating that ChEC-seq provides information on the directionality of TF-DNA interactions. Strikingly, similar DNA shape patterns are observed regardless of motif strength, indicating that the kinetics of ChEC-seq discriminates DNA recognition through sequence and/or shape. We propose that time-resolved ChEC-seq detects both high-affinity interactions of TFs with consensus motifs and sites preferentially sampled by TFs during diffusion and sliding.

## Introduction

Genome-wide determination of protein binding sites is of great interest for understanding normal and pathological cellular processes. Numerous techniques have been developed to map global protein-DNA interactions, and the most popular is formaldehyde crosslinking chromatin immunoprecipitation with high-throughput sequencing (X-ChIP-seq). Although ChIP-seq has been used to gain numerous insights into the regulation of DNA-templated processes, it has notable limitations attributable to crosslinking and sonication^1^. Formaldehyde, the most commonly used reagent for ChIP crosslinking, preferentially generates protein-protein crosslinks^2^ and can lead to epitope masking. X-ChIP-seq may also artificially inflate transient factor-chromatin interactions^3,4^, a problem that appears to be particularly acute at highly transcribed regions^5-7^. The resolution of X-ChIP-seq is also limited by sonication, though the addition of nuclease digestion steps (as in ChIP-exo, high-resolution X-ChIP, and ChIP-nexus) can greatly improve its resolution^8-10^. Additionally, sonication may introduce biases towards protein binding sites in open chromatin^11,12^. ChIP-seq performed without crosslinking, as in Occupied Regions of Genomes from Affinity-purified Naturally Isolated Chromatin (ORGANIC) profiling, circumvents issues associated with crosslinking and provides high resolution due to the use of micrococcal nuclease (MNase) to fragment chromatin^12,13^. However, the solubility of chromatin-associated proteins may be poor under the relatively gentle extraction conditions required for non-crosslinking methods^9^.

While ChIP-seq is most frequently used to map genome-wide protein-DNA interactions, a number of orthogonal methods, each involving fusion of chromatin proteins to DNA-modifying enzymes, have also been implemented. In one such method, DNA adenine methyltransferase identification (DamID)^14^, a protein of interest is fused to the Dam methyltransferase, resulting in methylation at regions bound by the protein and containing GATC sequences. In conjunction with microarray analysis, DamID has been used extensively to characterize genome-wide protein-DNA interactions in a range of model systems^15-19^. DamID allows the identification of protein binding sites in living cells without the need for crosslinking or immunoprecipitation, and, as it relies upon total DNA extraction rather than chromatin solubilization, it is quantitative and can thus be used with small amounts of starting material. However, the resolution of DamID is limited to kilobase-sized regions^15^ and the DNA-methylating activity of the fusion protein is constitutive. A second enzymatic method is Calling Card-seq, in which a chromatin protein of interest is fused to a transposase to facilitate targeted integration of transposons into the genome^20^. This method offers advantages similar to DamID with the added benefit of somewhat higher resolution, though it may be limited by transposase sequence preferences and also depends on the presence of restriction sites an appropriate distance from the inserted transposon to create templates for circularization and inverse PCR.

A third enzymatic method, chromatin endogenous cleavage (ChEC)^21^, makes use of a fusion protein comprising a chromatin protein of interest and MNase, which degrades unprotected DNA in the presence of calcium. ChEC has been used to characterize protein binding at specific genomic loci in yeast, such as the *GAL1-10* promoter, *HML,* and rDNA^21,22^, and has been used in conjunction with low-resolution microarray analysis to assess the association of nuclear pore components with the genome^23^. The benefits of ChEC are similar to those of DamID and Calling Card-seq, with resolution that is one to two orders of magnitude higher than that of these techniques, approaching base-pair resolution when analyzed by primer extension^23^. Importantly, ChEC is controllable, as robust MNase activity depends on the addition of calcium to millimolar concentrations, several orders of magnitude greater than the 50-300 nM free calcium observed in unstimulated yeast^24^ and mammalian cells^25-29^.

We asked whether a combination of ChEC with high-throughput sequencing (ChEC-seq) would allow high-resolution determination of protein binding sites on a genome-wide scale while circumventing issues with crosslinking, protein solubility, and antibody quality. lndeed, ChEC-seq yielded several times more binding sites for the budding yeast transcription factors (TFs) Abf1, Rap, and Reb1 than have been reported by ChIP-based methods. Making use of the inducible nature of ChEC, we found that binding sites for these TFs could be partitioned into two distinct temporal classes. The first displayed high levels of cleavage less than a minute after the addition of calcium and contained robust matches to known consensus motifs. In contrast, the second class of sites did not display appreciable levels of cleavage until several minutes after calcium addition and was depleted of motif matches. Sites containing motifs also displayed asymmetric cleavage patterns, indicating that ChEC-seq can detect directional TF-DNA binding. Strikingly, we found that sites both with and without motifs displayed notable DNA shape features relative to random sites, indicating that the kinetics of ChEC can separate TF binding sites (TFBSs) recognized by a combination of DNA shape and sequence or shape alone. We propose that rapidly cleaved sites, containing high-scoring motifs, represent direct, high-affinity binding of TFs to DNA, while slowly cleaved sites, with low-scoring motifs, are loci transiently sampled by TFs during diffusion and sliding due to their favorable shape profiles. Our results establish ChEC-seq as a robust genome-wide high resolution mapping technique orthogonal to ChIP-seq that we anticipate will be broadly applicable to numerous biological systems.

## Results

### Overview of the ChEC-seq experimental strategy

We generated a construct encoding a 3xFLAG epitope and MNase for PCR-based C-terminal tagging of endogenous loci in budding yeast. We chose to interrogate the genome-wide binding of the three canonical general regulatory factors: ARS Binding Factor 1 (Abf1), Repressor Activator Protein (Rap1), and RNA polymerase I Enhancer Binding protein (Reb1). Abf1 contains a bipartite DNA-binding domain (DBD) consisting of a zinc finger and an uncharacterized domain and regulates RNA polymerase II transcription as well as DNA replication^30^ and repair^31^. Rap1 contains a Myb-family helix-turn-helix DBD and regulates the expression of ribosomal protein genes^32^ and telomere length^33^. Reb1, like Rap1, contains a Myb-family helix-turn-helix DBD and is involved in the regulation of RNA polymerase I and II transcription^34-36^. ChEC in conjunction with southern blotting has been successfully used to map the binding of Reb1 to rDNA^37,38^. In addition, all three factors have been implicated in the formation of nucleosome-depleted regions (NDRs) at promoters throughout the yeast genome^39-41^.

TFs are often expressed at levels expected to drive nonspecific interactions with chromatin via mass action^42,43^ and scan for their binding sites via trial-and-error sampling of sites on chromatin^4^. We therefore anticipated that a substantial fraction of cleavages in the TF-MNase strains could be due to random diffusion and collision of the fusion proteins with chromatin. To control for this, we generated a strain harboring a construct encoding 3xFLAG-tagged MNase fused to an SV40 nuclear localization signal under the control of the *REB1* promoter integrated at the *URA3* locus (‘free MNase’). As there are more molecules of Reb1 than either Abf1 or Rap1 in a yeast cell^44^, we surmised that free MNase driven by the Reb1 promoter would also serve as a suitable control for Abf1 and Rap1 ChEC-seq experiments. The free MNase control is analogous to the unfused Dam control used in DamID experiments^15^. Expression of free MNase and TF-MNase fusions was well tolerated, as cells displayed no overt growth phenotype (Figure 1a), though they showed increased background DNA damage as assessed by yH2A levels (Figure 1b), in the absence of exogenous calcium.

**Figure 1.**
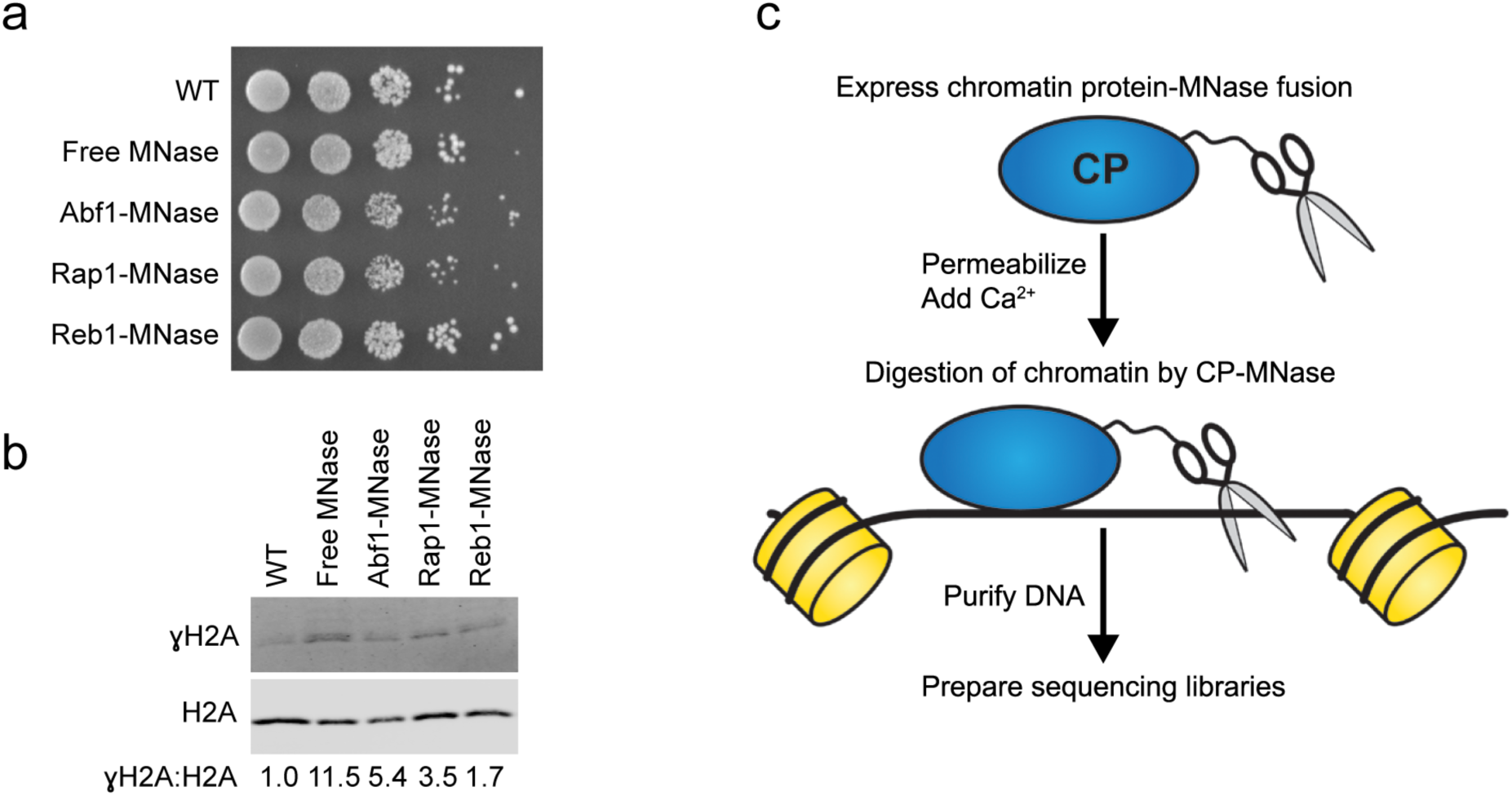
Phenotypic characterization of strains bearing MNase-tagged TFs. (**a**) Growth of free MNase and TF-MNase strains on YPD. (**b**) Western blot analysis of H2A serine 129 phosphorylation (ɣH2A) in free MNase and TF-MNase strains. The ɣH2A/total H2A ratio is indicated under each lane. (**c**) ChEC-seq workflow. Yeast cells expressing a chromatin protein (CP) of interest genetically fused to MNase are permeabilized with digitonin and calcium is added to induce cleavage by CP-MNase. Digested DNA is then purified and prepared for high-throughput sequencing.

We followed the previously described *in vivo* ChEC protocol^21^, wherein living yeast cells are permeabilized with digitonin prior to the addition of Ca^2+^ to induce chromatin cleavage (Figure 1c). We presumed that treatment of permeabilized cells with Ca^2+^ would generate both specific cleavages at TFBSs and nonspecific cleavages resulting from mass action-driven interactions of the TF-MNase fusions with chromatin, leading to the generation of small protected fragments representing TFBSs. We thus performed size selection of ChEC DNA prior to sequencing library preparation to enrich for small DNA fragments.

Prior to size selection, we analyzed the kinetics of bulk genomic DNA degradation by TF-MNase fusions and free MNase by agarose gel electrophoresis. Analysis of widely-spaced, minute-scale time points revealed notable smearing of genomic DNA by all three TF-MNase fusions by the 2.5 m time point. In contrast, 2.5 m of digestion in the free MNase strain yielded only very slight smearing of high molecular weight genomic DNA (Supplementary Figure 1a). This pattern persisted until 20 m, when robust smearing of genomic DNA by free MNase could be observed. We interpret these patterns to indicate that specific cleavage mediated by TF-MNase fusions occurs relatively rapidly, at specific sites, following Ca^2+^ addition, while digestion with free MNase takes longer due to the fact that it is not specifically targeted to any sites on chromatin. A similar pattern was observed with second-scale digestion of DNA with Reb1-MNase, where slight smearing of genomic DNA was evident as early as 10 s, but no such degradation was observed in the free MNase strain (Supplementary Figure 1b).

### ChEC-seq maps genome-wide TF binding

Visualization of mapped ChEC-seq fragments revealed robust, discrete sites of cleavage over negligible background signal with variable dynamic range depending on the extent of digestion (Figure 2a-c). A modest amount of cleavage was observed as quickly as 10 s, presumably due to activation of TF-MNase molecules bound to DNA at the outset of the time series, and increased markedly by 20 s. Signal-to-noise was generally highest at the 30 and 40 s time points, and dynamic range decreased thereafter, presumably due to TF unbinding and subsequent digestion of binding sites combined with an overall increase in background signal due to increasing nonspecific cleavage. Strong cleavage was observed at Abf1 and Reb1 sites identified by ORGANIC profiling^12^ (Figure 2a,c) and a Rap1 site identified by ChIP-exo^8^ (Figure 2b). Free MNase digestion yielded only nonspecific patterns of cleavage with a much reduced dynamic range compared to that seen for Abf1, Rap1, and Reb1 at the same genomic regions (Figure 2a-c), indicating that the patterns of cleavage in the TF-MNase strains are specifically due to chromatin targeting of MNase by fusion to TFs and that ChEC-seq is unlikely to suffer from a chromatin accessibility bias. We also assessed the specificity of our ChEC-seq data by determining ChEC cleavages at Abf1, Rap1, and Reb1 peaks previously determined by various ChIP methods^8,12,45^ and at an identical number of randomly generated genomic regions. Average enrichment of cleavages within ChIP peaks was 8.5-fold to 70-fold higher than at random sites (Supplementary Figure 1c-e), further indicating that ChEC-seq specifically detects TFBSs.

**Figure 2.**
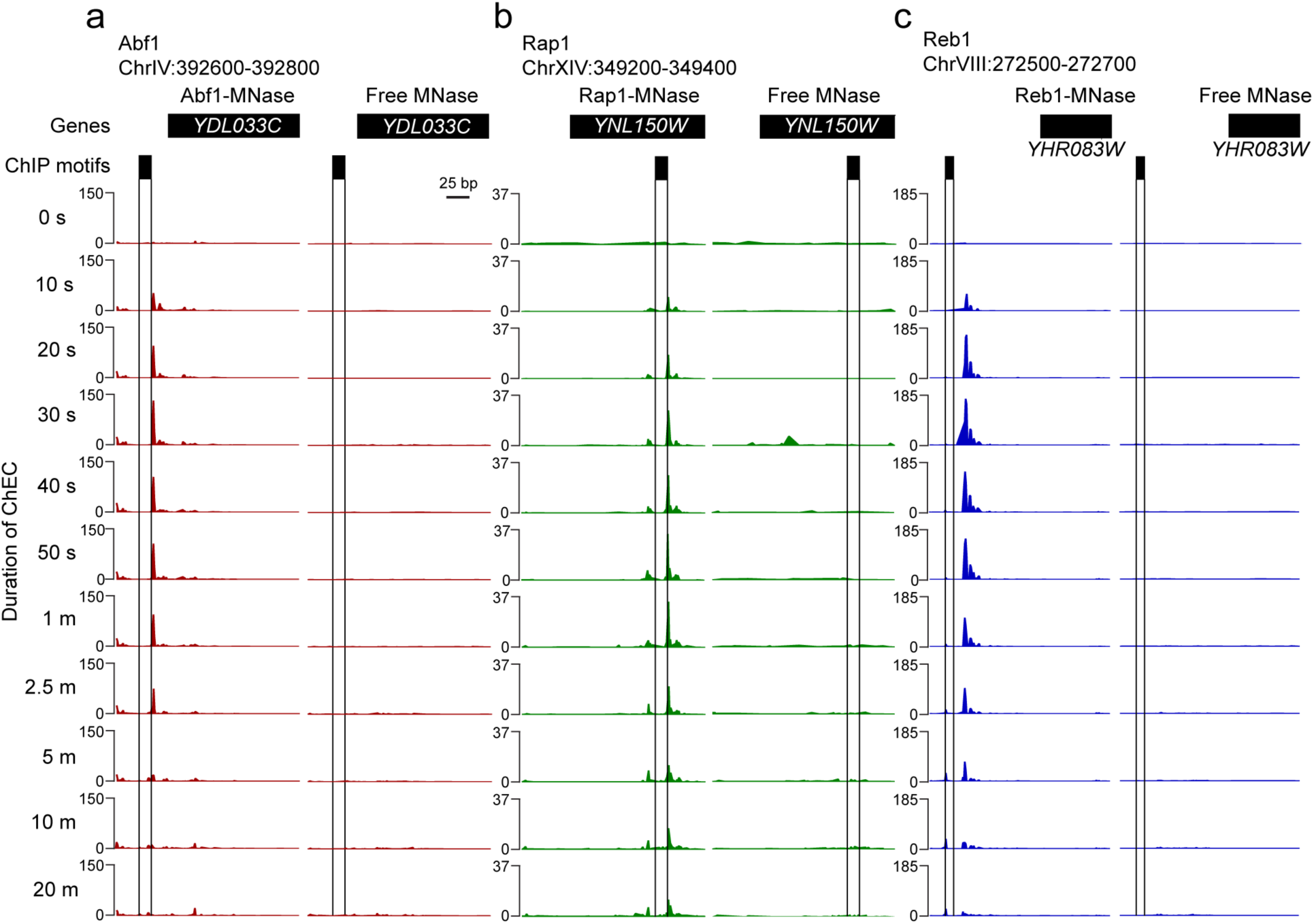
Genome-wide mapping of TF binding with ChEC-seq. Tracks of ChEC-seq signal for (**a**) Abf1, (**b**) Rap1, and (**c**) Reb1 along the indicated 200 bp segments of the genome. The positions of Abf1 and Reb1 motifs detected within ORGANIC peaks and a Rap1 motif detected within a ChIP-exo peak is indicated. Tracks of free MNase ChEC-seq signal at these genomic regions are also shown.

### ChEC-seq reveals temporally distinct classes of TFBSs

Data generated by steady-state methods such as ChIP and DamID have two dimensions: binding site location and occupancy. In addition to these, ChEC-seq data provides a third dimension: time. We wondered if analysis of ChEC-seq cleavage kinetics could provide insight into the modes of DNA recognition by TFs. We thus determined the maximum signal for each base position in the genome irrespective of digestion time and called peaks on this composite dataset using a genome-wide thresholding approach, followed by analysis of signal within each peak across all time points.

For Abf1, we detected 12,351 peaks (Supplementary Data 1). Of note, this is an order of magnitude more peaks than was detected using ORGANIC profiling^12^. To ask if there are temporally distinct classes of Abf1 sites, we performed unsupervised *k*-means clustering of peak signal with *k* = 2. This analysis revealed a striking partitioning of the data into fast and slow classes based on the time point at which maximum signal was reached (Figure 3a, Supplementary Figure 2a). We wondered if this temporal partitioning of sites might reflect differences in the affinity of Abf1 for the DNA sequences underlying these peaks and so scored each site using a previously published position frequency matrix (PFM)^45^. We observed a robust relationship between temporal class and motif strength, with the highest motif scores corresponding to the fast class and lower motif scores corresponding to slow sites (Figure 3a). Consistent with this, *de novo* motif discovery within fast sites revealed a robust match to the Abf1 consensus, similar to that previously determined by ORGANIC^12^, while a nonspecific AT-rich sequence was the most enriched sequence in slow sites (Figure 3a). To test the reproducibility of these data, we performed two additional replicates at the 30 s time point and compared the sum of cleavages within each peak. This comparison revealed excellent correspondence between replicates (Spearman’s rank correlation *ρ* = 0.966-0.981; Supplementary Figure 3a). Extending these analyses to Rap1 and Reb1, we detected 7,260 Rap1 peaks, over twelve times the number of Rap1 peaks previously determined by ChIP-exo^8^, and 8,268 Reb1 peaks, greater than four times the number of Reb1 sites previously reported in ChIP studies^8,12^ (Supplementary Data 1). As was observed for Abf1 peaks, Rap1 (Figure 3b, Supplementary Figure 3b) and Reb1 (Figure 3c, Supplementary Figure 3c) peaks could be divided into fast and slow clusters also distinguished by motif strength. *De novo* motif discovery using fast but not slow Rap1 and Reb1 peaks also revealed robust matches to previously determined consensus sequences, similar to those found by ChIP-exo and/or ORGANIC, and AT-rich motifs bearing some resemblance to those found in slow Abf1 sites were found in Rap1 and Reb1 slow sites (Figure 3b-c). Cleavage levels for Rap1 and Reb1 peaks were also robust across 30 s replicates (Rap1 Spearman’s rank correlation *ρ* = 0.963-0.990; Supplementary Figure 3b and Reb1 Spearman’s rank correlation *ρ* = 0.945-0.989; Supplementary Figure 3c). We also analyzed the reproducibility of cleavage levels across 2.5 m replicates for Abf1 and found robust correlations (Spearman’s rank correlation *ρ* = 0.901-0.958; Supplementary Figure 3d).

**Figure 3.**
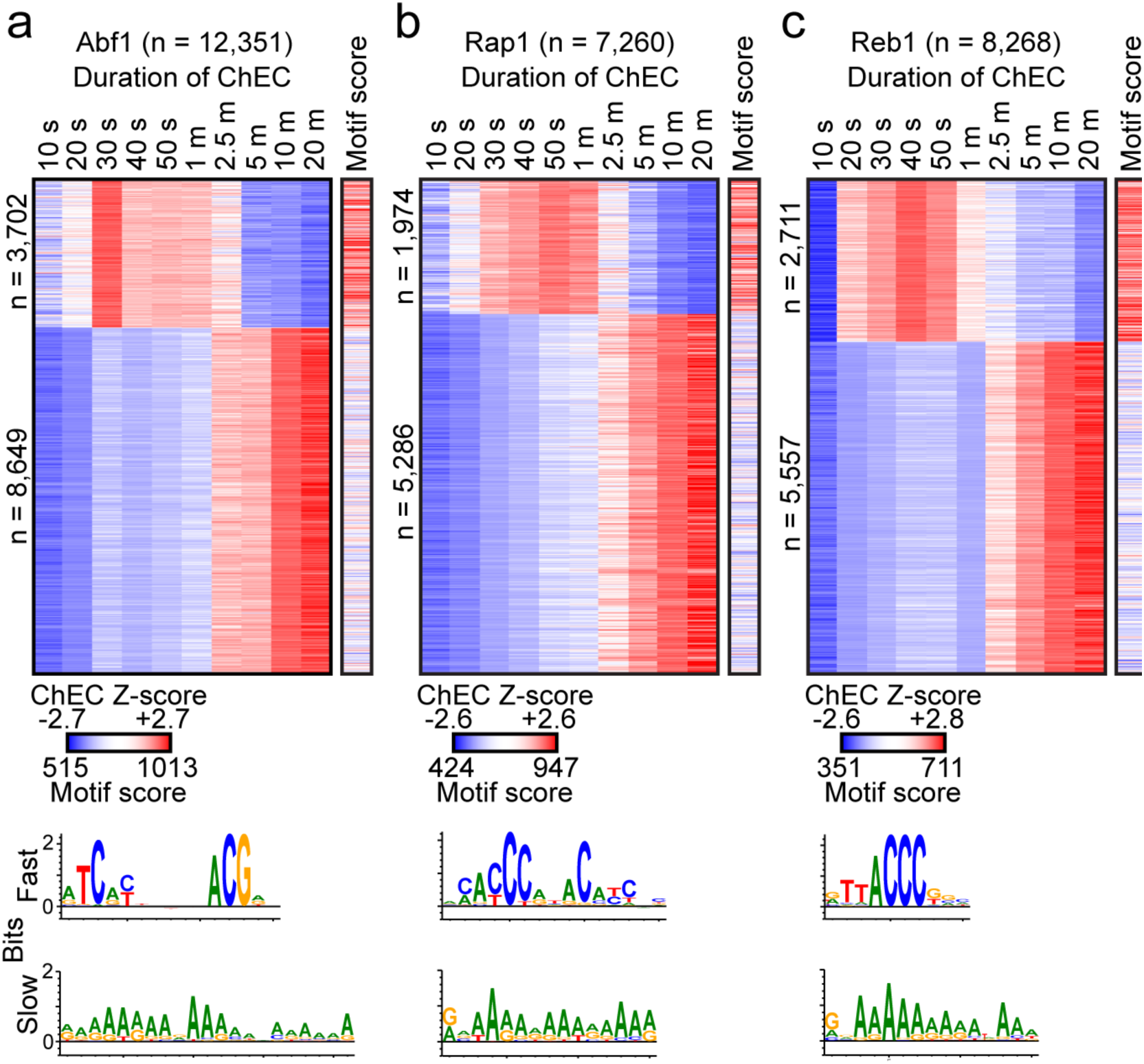
Temporal analysis of ChEC-seq data reveals distinct classes of TFBSs. Heatmaps of raw Z-scored, clustered ChEC-seq signals and motif scores for (**a**) Abf1, (**b**) Rap1, and (**c**) Reb1 sites. The sequence logos of the highest-scoring motif discovered by MEME-ChIP and plotted with LogOddsLogo in fast and slow sites are displayed below the heatmap for the corresponding factor.

We considered the possibility that, because slow sites generally lack robust consensus motifs, these sites could simply be due to chromatin accessibility or other biases. If this were the case, we would expect the majority of slow sites to overlap across datasets for multiple TFs. However, analysis of the overlap of slow sites between TFs revealed that 7,550/8,649 (87.3%) of Abf1 slow sites, 4,115/5,286 (77.8%) of Rap1 slow sites, and 4,974/5,557 (89.5%) of Reb1 slow sites were unique. This suggests that slow sites represent preferred sites without robust consensus motifs for TFs that may be sampled during diffusion and sliding^4^.

To further assess the specificity of our ChEC-seq peaks, we called peaks on the free MNase dataset (Supplementary Data 1) and used these peaks to generate a false discovery rate (FDR) for each TF-MNase peak set. The FDR was defined as the percentage of TF-MNase peaks that were also found in the free MNase dataset. FDRs calculated were 2.37% for Abf1, 3.71% for Rap1, and 2.42% for Reb1. These small FDRs indicate that at most a very minor fraction of the peaks in each experiment (Abf1, 292/12,353; Rap1, 269/7,260; Reb1, 200/8,268) might be artifactual. These FDRs may be overestimates, because a number of free MNase peaks occur in telomeric and rDNA regions, which tend to have high signal in ChIP experiments, but to which Rap1 and Reb1 have been shown to bind by multiple ChIP approaches^8,12^. The correlation between TF and free MNase signals at peaks was also very poor (Spearman’s rank correlation *ρ* = 0.080-0.118; Supplementary Figure 4). We conclude from these analyses that ChEC-seq peaks are dependent on targeting of MNase to chromatin by fusion to a TF.

**Figure 4.**
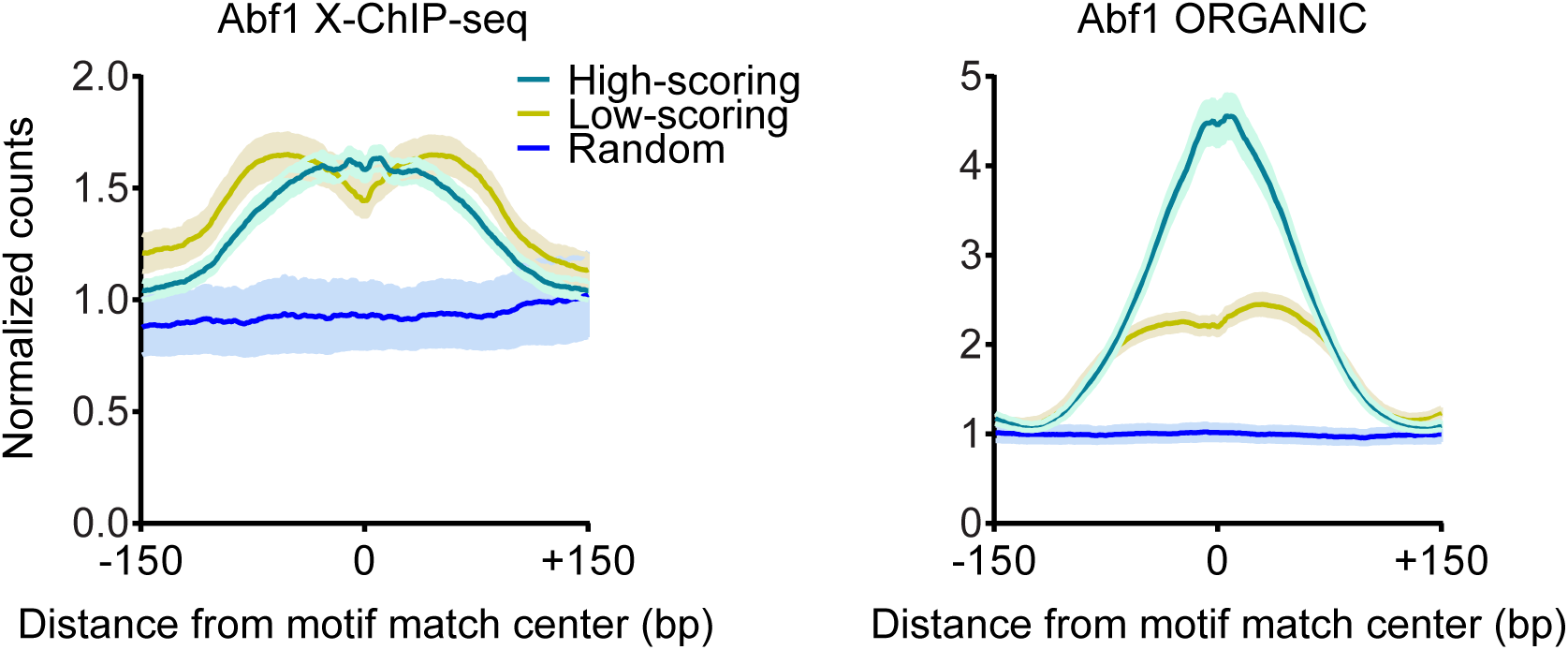
ChEC-seq peaks display enrichment in ChIP experiments. Average plots of Abf1 X-ChIP-seq and ORGANIC signal around high-scoring, low-scoring, and random sites. Shaded areas represent 95% confidence intervals.

### ChEC-seq peaks are enriched in ChIP-seq datasets

Recent work has indicated that the extended formaldehyde crosslinking (10-15 min) usually performed in X-ChIP experiments captures transient interactions of TFs with degenerate motifs during binding site scanning^3,4^. We thus anticipated that X-ChIP experiments would capture ChEC-seq sites regardless of their temporal profile and motif strength. As our cluster analysis revealed two distinct classes of sites distinguishable by motif strength, we refined the binary classification of ChEC-seq sites by parsing sites by motif strength for further analysis. For the purposes of all subsequent analyses, we define sites with a motif match p-value < 0.001 as ‘high-scoring sites’ and those with a motif match p-value ≥ 0.001 as ‘low-scoring sites’.

While X-ChIP methods capture protein-DNA interactions regardless of duration when a long crosslinking step is used, ORGANIC preferentially enriches for stable, high-affinity protein-DNA interactions^12^. We thus hypothesized that, if low-scoring sites were representative of transient chromatin interactions during scanning, they would be less enriched relative to high-scoring sites in ORGANIC compared to X-ChIP experiments. To examine this, we performed an X-ChIP experiment in which chromatin was digested with MNase (MNase-X-ChIP-seq) for Abf1. We then generated average plots of Abf1 X-ChIP-seq and ORGANIC signal around each class of sites. As we hypothesized, Abf1 enrichment was essentially equal at high- and low-scoring sites when assessed by X-ChIP-seq (Figure 4). In contrast, average Abf1 enrichment was ~2- fold lower at low-scoring compared to high-scoring sites when assayed by ORGANIC (Figure 4). These analyses further support the specificity of ChEC-seq and suggest that low-scoring ChEC-seq sites represent transient interactions of TFs with preferred sites during binding site scanning.

We next assessed the overlap of ChEC-seq peaks with peaks obtained by various ChIP approaches. High-scoring Abf1 ChEC-seq peaks overlapped with 929/1,277 (72.7%) of peaks discovered by X-ChIP-chip^45^ and/or OrGAnIC^12^, while low-scoring Abf1 ChEC-seq peaks intersected with 696/1,277 (54.5%) of these ChIP peaks (Supplementary Figure 5a). High-scoring Rap1 ChEC-seq peaks overlapped with 395/654 (58.9%) of peaks reported by X-ChIP-chip and/or ChIP-exo, while low-scoring Rap1 ChEC-seq peaks coincided with 270/654 (41.3%) of these ChIP peaks (Supplementary Figure 5b). High-scoring Reb1 ChEC-seq peaks overlapped with 1,839/2,820 (65.2%) of peaks discovered by X-ChIP-chip and/or ORGANIC, while low-scoring Reb1 ChEC-seq peaks corresponded to 1,136/2,820 (40.3%) of these ChIP peaks (Supplementary Figure 5c). These results indicate that ChEC-seq captures a substantial fraction of TFBSs previously identified by ChIP-based methods. Furthermore, the stronger overlap of high-scoring ChEC-seq peaks with ChIP peaks, most of which were reported to contain consensus motifs, is consistent with the robust motif matches in this class of ChEC-seq sites.

**Figure 5.**
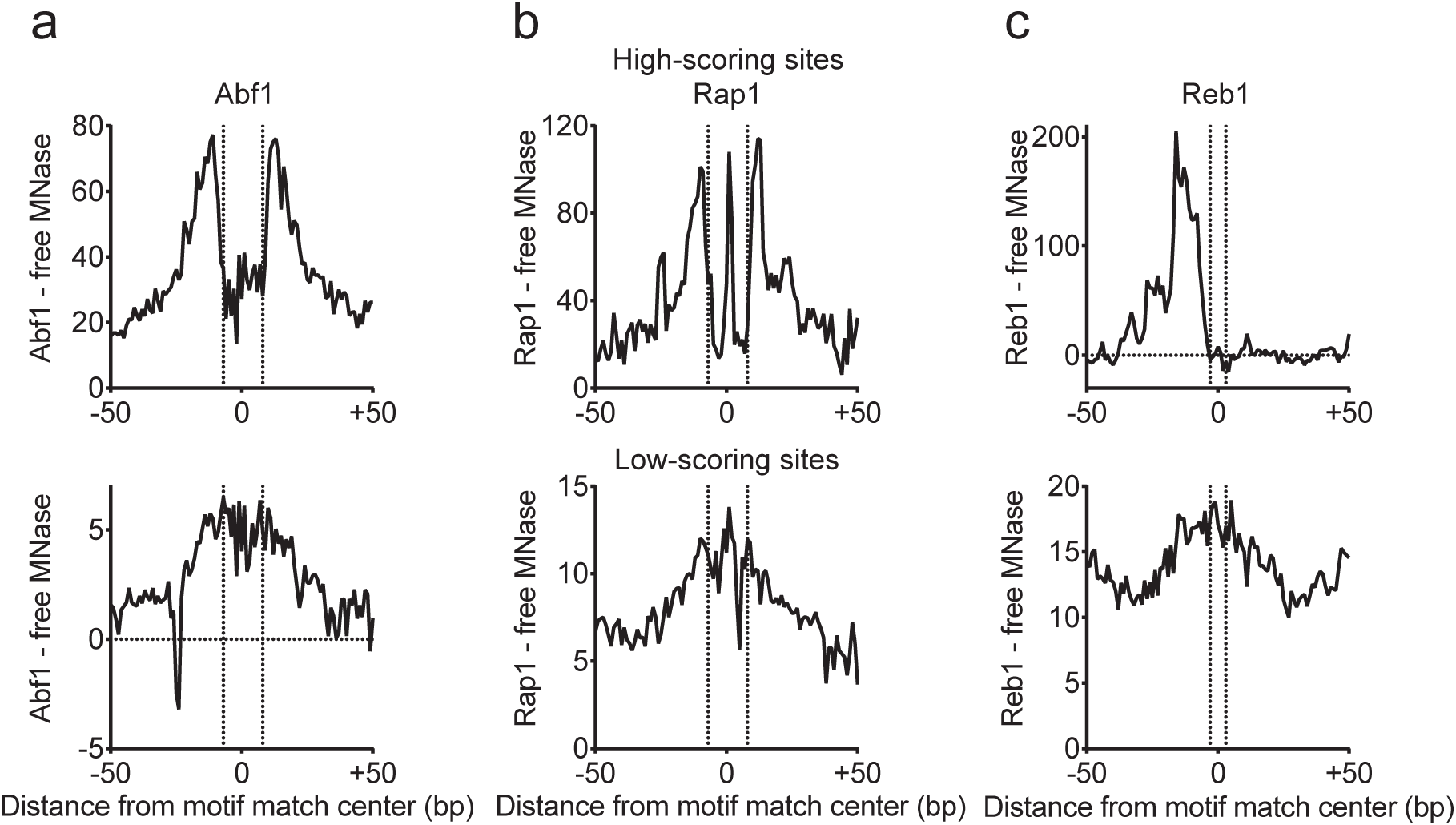
ChEC-seq sites display distinct cleavage profiles based on motif strength. (**a**) Average plots of Abf1 cleavage around high-scoring and low-scoring Abf1 sites. The boundaries of the motif match are indicated by vertical dotted lines. (**b**) Same as (**a**) but for Rap1 sites. (**c**) Same as (**a**) but for Reb1 sites. Sites are oriented such that the best match to the consensus motif used for scoring is left to right on the top strand. The *y*-axis scale has been expanded in the lower panels to reveal details in the low-scoring site profiles.

### ChEC-seq detects directionality in protein-DNA interactions

As we fused MNase to the C-terminus of each TF, we wondered if we could obtain structural information about the orientation of each TF on DNA and thus analyzed cleavage patterns around high-scoring and low-scoring sites for each TF. High-scoring Abf1 sites showed robust and essentially equivalent peaks of cleavage upstream and downstream of the motif match (Figure 5a). We also observed a moderate frequency of cleavage within the motif match, likely indicating MNase accessibility of the nonspecific spacer between the two specific halves of the Abf1 motif^46^. Low-scoring Abf1 sites displayed low levels of cleavage throughout the motif match and the upstream and downstream regions (Figure 5a). High-scoring Rap1 sites also showed strong cleavage peaks to either side of the motif match, as well as a strong cleavage peak in the center of the motif match (Figure 5b). The cleavage in the center of the Rap1 motif match may be attributable to cleavage within the 3 bp linker that spans the two hemi-sites making up the Rap1 consensus sequence^47^. Low-scoring Rap1 sites displayed a noisier version of the tripartite cleavage pattern observed at more robust motif matches (Figure 5b). In contrast to Abf1 and Rap1, high-scoring Reb1 sites displayed a strongly asymmetric pattern of cleavage, with nearly all cleavages occurring upstream of the motif match (Figure 5c). However, this pattern was not observed at low-scoring Reb1 sites (Figure 5c). These data suggest that ChEC-seq, with appropriate structural consideration of the protein under study, is capable of providing information about the orientation of TFs bound to DNA.

Though all three TFs tested presumably bind in a directional manner to their nonpalindromic motifs, robust cleavage asymmetry was observed only in Reb1 experiments (Figure 5c). This may be explained by structural flexibility of the proteins under study and/or the relatively long length of the TF-MNase linker (33 aa). In the case of Abf1, its C-terminal ~200 aa contain no domains and as such might be quite flexible, allowing MNase on a long linker to cleave on either side of its binding sites. Rap1 contains a C-terminal RCT protein-protein interaction domain, but this domain is predicted to be positioned above the central Rap1 DBDs and so may afford MNase access to DNA on either side of its binding sites^48^.

To investigate the effect of linker length on the observed ChEC-seq profiles, we shortened the TF-MNase linker from 33 to 8 aa (short linker strains are hereafter referred to as TF-SL) and performed ChEC-seq. The Abf1-SL strain displayed the same cleavage patterns at Abf1 sites determined in the longer linker strain, suggesting that it is structural flexibility in the Abf1 C-terminus that dictates cleavage patterns in these strains (Figure 6a). As expected, the Reb1-SL strain displayed a cleavage pattern very similar to that of the Reb1 strain (Figure 6b). We did not observe cleavage in the Rap1-SL strain, potentially due to the long expected distance of the Rap1 C-terminus from DNA^48^.

**Figure 6.**
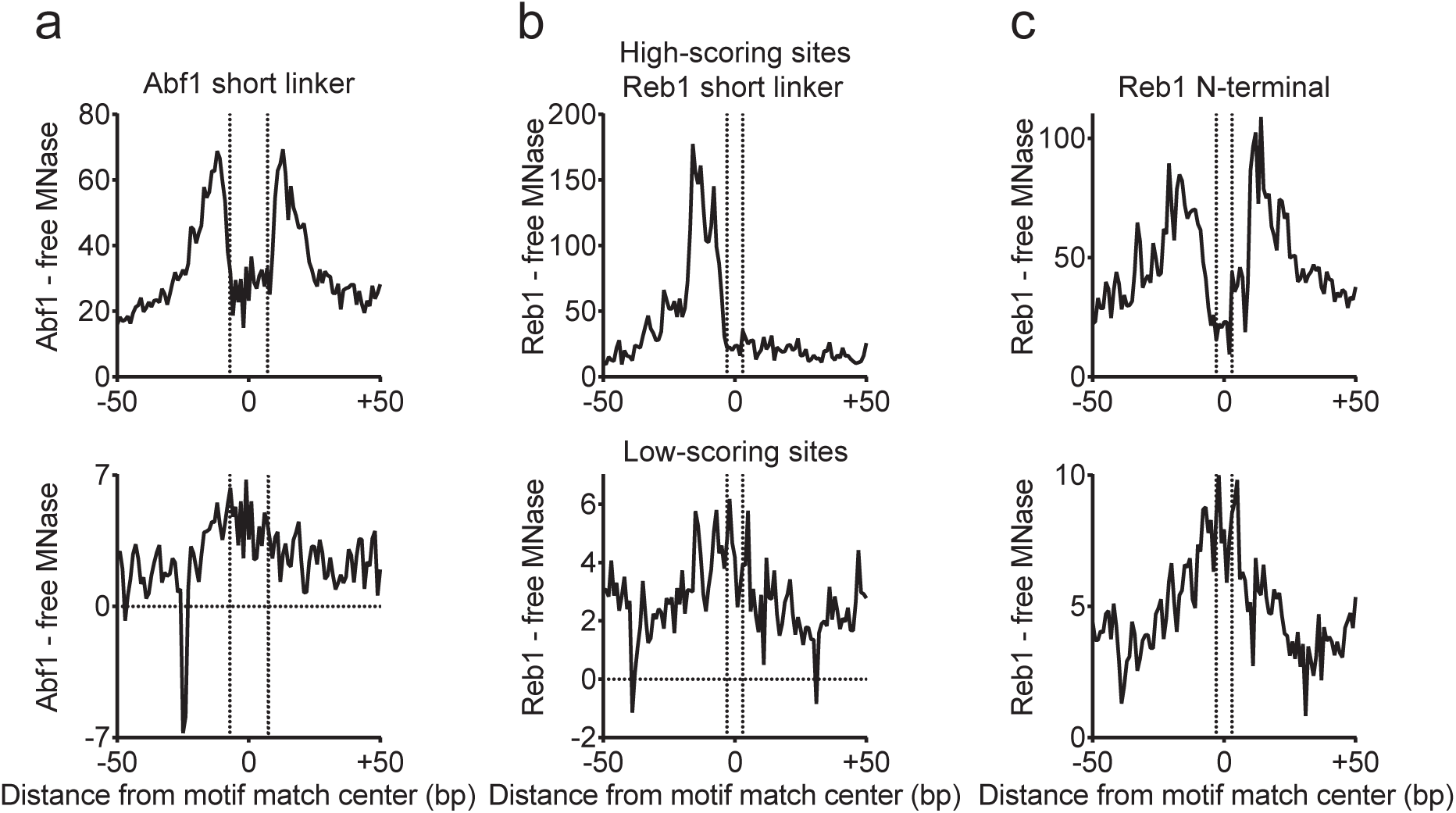
TF-MNase linker alteration modulates cleavage patterns. **a**) Average plots of Abf1-SL cleavage around high-scoring and low-scoring Abf1 sites. The boundaries of the motif match are indicated by vertical dotted lines. (**b**) Same as (**a**) but for Reb1 sites. (**c**) Average plots of Reb1 cleavage around high-scoring and low-scoring Reb1 sites. The *y*-axis scale has been expanded in the lower panels.

To further explore the effects of structural flexibility on ChEC-seq cleavage patterns, we expressed Reb1 tagged with MNase at its N-terminus from a plasmid and performed ChEC-seq. Tagging of the Reb1 N-terminus with MNase resulted in a symmetric cleavage pattern (Figure 5c) similar to that observed in the Abf1 and Rap1 strains (Figure 5a,b), suggesting that structural flexibility of the protein under study has a substantial effect on the cleavage pattern observed.

### High-scoring and low-scoring TFBSs share DNA shape features

What drives preferential interaction of TFs with low-scoring sites? Recent work has suggested that, in addition to sequence, DNA shape can drive recognition of specific loci by TFs^49,50^. We thus wondered if low-scoring sites might contain DNA shape features conducive to recognition and so analyzed minor groove width (MGW), Roll, propeller twist (ProT), and helix twist (HelT) at high-scoring and low-scoring sites for each TF. For each TF and category of DNA shape features, we compared the patterns between high-scoring and low-scoring sites. Pearson correlation coefficients (PCCs) close to 1 and large Kolmogorov-Smirnov (KS) *p* values indicate similar DNA shape profiles. Abf1 high-scoring and low-scoring sites displayed highly similar patterns of MGW (PCC = 0.98; KS p = 0.68), Roll (PCC = 0.99; KS p = 0.68), ProT (PCC = 0.98; KS p = 0.94), and HelT (PCC = 0.98; KS p = 0.68) (Figure 7a). Similarly, Rap1 high-scoring and low-scoring sites showed very similar MGW (PCC = 0.97; KS p = 0.18), Roll (PCC = 0.98; KS p = 0.68), ProT (PCC = 0.98; KS p = 0.99), and HelT (PCC = 0.98; KS p = 0.94) (Figure 7b). Lastly, analysis of Reb1 high-scoring and low-scoring sites likewise yielded good correspondence of MGW (PCC = 0.94; KS p = 0.83), Roll (PCC = 0.98; KS p = 0.83), ProT (PCC = 0.99; KS p = 0.83), and HelT (PCC = 0.93; KS p = 0.99) (Figure 7c). Shape parameters at randomly selected binding sites displayed weak correlation with and significant difference (PCC < 0.3; KS p < 0.05) from those at both high-scoring and low-scoring sites for all three factors.

**Figure 7.**
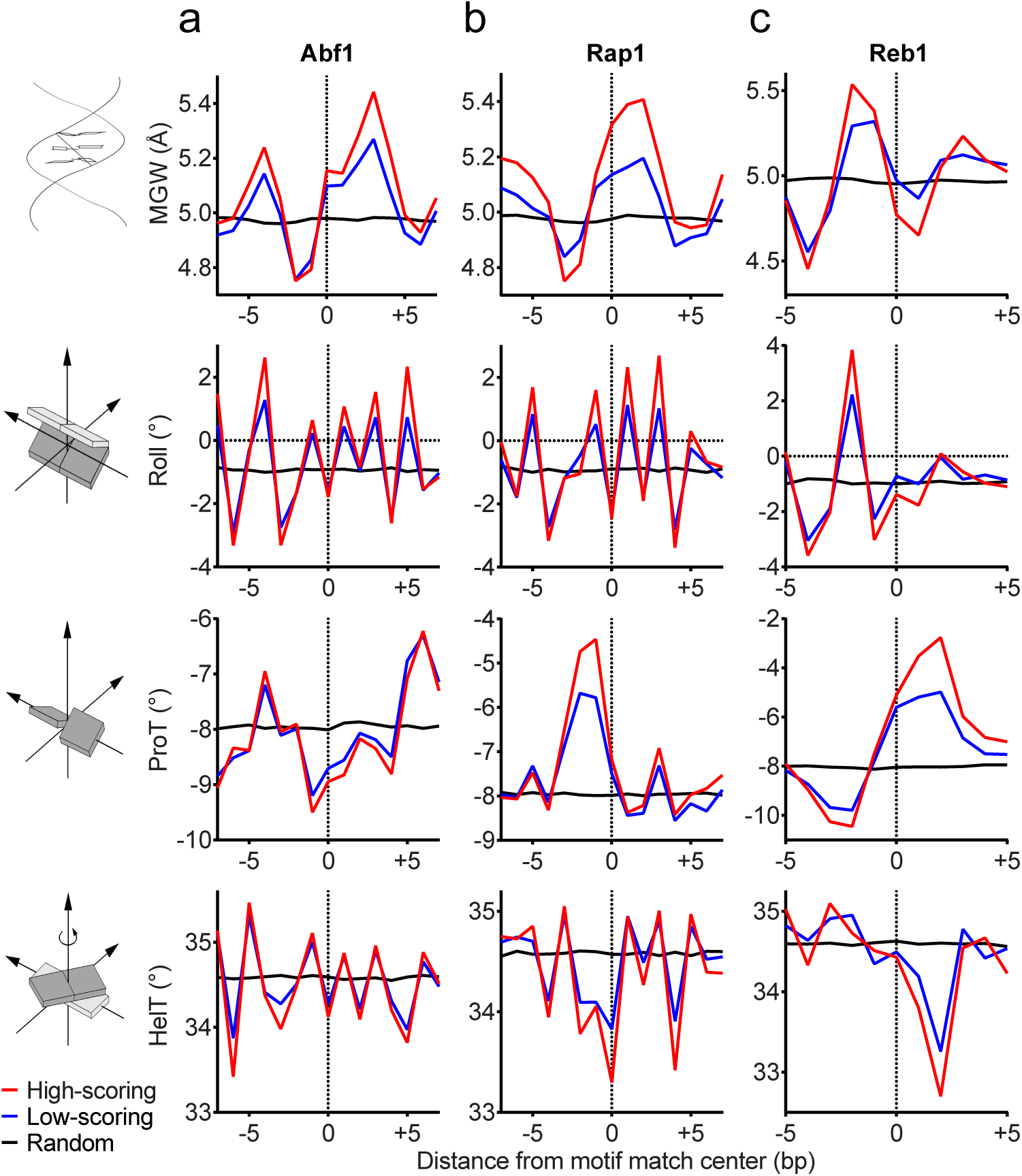
ChEC-seq-derived sites display distinctive DNA shape profiles for each TF regardless of the strength of a consensus motif. Average profiles of minor groove width (MGW), Roll, propeller twist (ProT), and helix twist (HeIT) at high-scoring, low-scoring, and random (**a**) Abf1, (**b**) Rap1, and (**c**) Reb1 sites. Plots were centered on the middle of the best match to the consensus motif used for scoring within each peak. Schematic representations of shape features are shown to the left of the row of plots for the corresponding feature.

We next asked if a small subset of low-scoring sites with relatively good matches to the consensus might be driving the DNA shape trends observed in the average plots. To test this, we generated heatmaps for each shape parameter and TF ranked by motif match p-value. Heatmaps for Abf1 (Supplementary Figure 6), Rap1 (Supplementary Figure 7), and Reb1 (Supplementary Figure 8) high-scoring and low-scoring sites were highly similar, suggesting that the identified shape patterns are not dependent on the presence of a strong consensus motif. We also compared the relative abilities of a shape model and a sequence model to distinguish between sites containing high-scoring and low-scoring motifs using L2-regularized multiple linear regression. The resulting values for area under the receiver operating characteristic (AUROC) showed that sequence is a better discriminator between high-scoring and low-scoring sites than shape (Supplementary Figure 9), indicating that DNA shape is more similar than sequence between these two classes of sites. These results suggest that ChEC-seq kinetics separates TFBSs on the basis of their recognition mode (sequence and shape versus shape alone).

**Figure 8.**
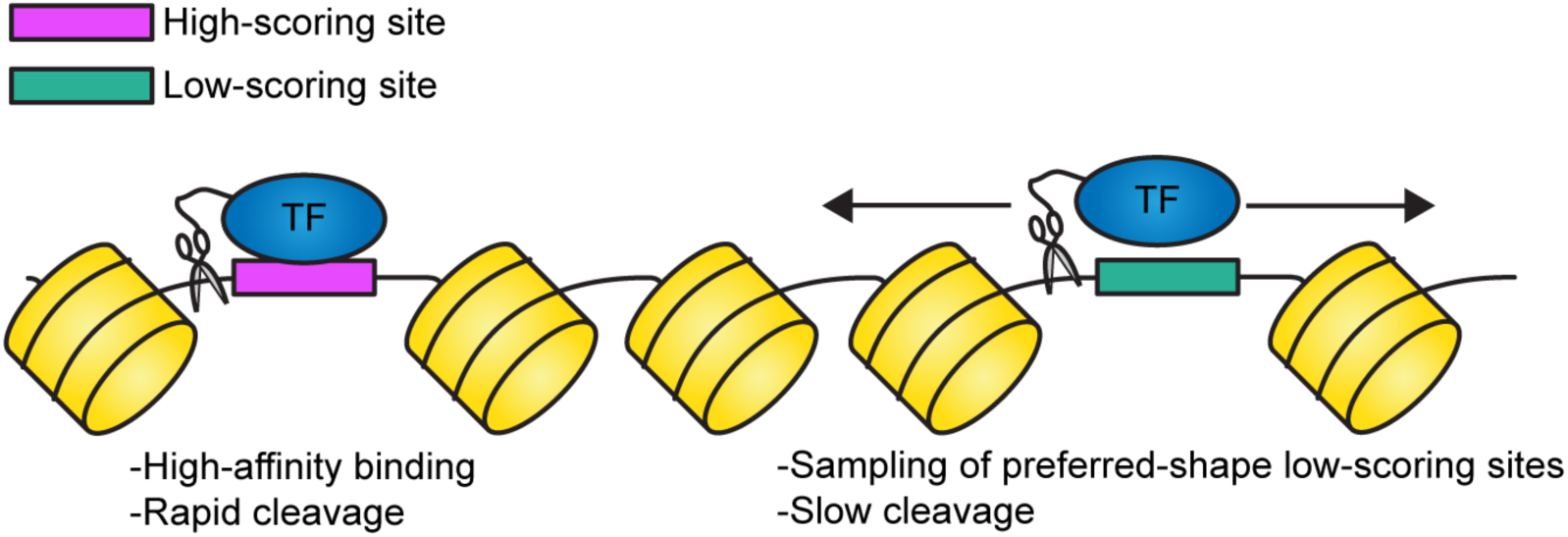
Characteristics of high-scoring and low-scoring ChEC-seq sites. A schematic representation of two TFBSs with differing affinity for the cognate TF. On the left, at a high-scoring site, a TF binds its cognate motif with high affinity, likely forming hydrogen bonds and other base-specific contacts, thus rapidly generating high levels of cleavage at specific sites. On the right, at a low-scoring site, a tF scans along the genome by sliding and sampling shape-preferred low-scoring sites, likely without forming contacts with the functional groups of the bases, thus generating dispersed low-level cleavages around these sites.

## Discussion

We have shown that ChEC-seq robustly identifies global protein-DNA interactions with high spatial and temporal resolution. Time-resolved analysis of ChEC-seq sites distinguishes two classes of TFBSs by cleavage kinetics and motif strength. High-scoring sites exhibited rapid cleavage during ChEC time courses, suggesting that they are already partially occupied when calcium is added and rapidly reach maximum occupancy. In contrast, low-scoring sites reached cleavage maxima twenty minutes after calcium addition but were largely unique to each TF, arguing that they do not simply reflect DNA accessibility *in vivo.* Strikingly, high-scoring and low-scoring sites displayed very similar profiles of DNA shape features.

The fact that low-scoring sites are only robustly detected several minutes into the ChEC time course suggests two possibilities: 1) low-scoring sites are only robustly cleaved after high-scoring sites are digested away or 2) the transient nature of TF-DNA interactions at low-scoring sites necessitates several minutes of incubation to detect robust cleavage levels. Although our current results cannot distinguish these two possibilities, the ability of ChEC-seq to kinetically separate high-affinity binding events and transient sampling interactions is a distinct advantage of the method over existing mapping approaches. We speculate that high-scoring sites represent sites of high-affinity protein-DNA interactions driven by recognition of both DNA sequence and shape, while low-scoring sites are lower affinity sites preferentially sampled during binding site scanning due to their favorable shape profiles (Figure 8). Our observations suggest that TFs first physically recognize sites with similar shapes and then further narrow down that group through the formation of sequence-specific hydrogen bonds with motifs. In this regard, DNA shape could serve as a trigger for a shift in protein conformation between the previously proposed scanning and folding modes^51^, which facilitate rapid binding site searching and stable protein-DNA interactions, respectively. Thus, it may be that DNA shape and protein conformation work in concert to limit the binding site search space.

A common finding from large-scale mapping efforts is that many TFs with well-established sequence specificities bind more sites without than with consensus motifs. For instance, for a group of 36 TFs with well-known DNA binding specificities, ~36-100% of X-ChIP-seq sites reported by the ENCODE project do not contain consensus motifs^52^. It has been proposed based on these and other similar results that indirect tethering of proteins to DNA via protein-protein interactions is more prevalent than is generally thought. Alternatively, these sites might represent transient interactions with DNA that are captured and inflated by cross-linking^53^. Consistent with this latter interpretation, analysis of Sox2 association with the genome by live-cell microscopy showed that its residence time at sites with poor consensus motifs is ~15-fold shorter than at sites with strong motifs, despite the fact that these sites can be captured in ChIP-exo experiments^4^. Notably, these transient interactions appear to be quite prominent, as Sox2 undergoes nearly 100 diffusions prior to locating a consensus binding site. Our results are consistent with this idea, indicating that the majority of sites without motif matches represent capture of transient scanning interactions with sites displaying preferred DNA shape features. ChEC-seq is thus a powerful tool for distinguishing high-affinity, sequence-dependent interactions from interactions with preferred low-scoring sites during sampling of binding sites *in vivo*, which is not possible with current steady-state mapping technologies. The high spatiotemporal resolution of ChEC-seq allowed us to gain mechanistic insights into TF-DNA sliding, which will likely enable the development of models for the search of TFBSs under *in vivo* conditions.

ChEC-seq has notable advantages relative to genome-wide mapping methodologies based on ChIP or other enzymatic fusions. In particular, the inducible nature of ChEC-seq provides kinetic information not currently available on a genome-wide scale, and it was the rapidly inducible nature of ChEC-seq that allowed us to separate TFBSs based on their recognition by DNA sequence and shape or shape alone. ChEC-seq also revealed structural information on TFs bound to their cognate sites. One limitation of ChEC-seq is that multiple time points must be performed to capture fast and slow sites; however, one early (i.e. 30 s) and one late (i.e. 20 m) should be sufficient to capture both classes. The major limitation of ChEC-seq is the requirement for expression of a fusion protein. Although tagging of endogenous loci has generally been laborious in non-yeast systems, the recent development of CRISPR/Cas9-based genome editing^54^ has greatly simplified tagging, and we speculate that MNase tagging of endogenous loci in non-yeast systems will be commonplace to generate reagents for genome-wide mapping of protein binding with high spatial and temporal resolution.

## Methods

### Yeast strain construction

The yeast ChEC-tagging vector pGZ108 was constructed by insertion of a PCR amplicon encoding a 3xFLAG tag and aa 83-231 of MNase (representing the mature sequence of MNase, GenBank P00644) followed by two stop codons (TAGTAG) between the PacI and AscI sites of pFA6a-3HA-kanMX6^55^ (Addgene #39295), replacing the 3xHA tag. The total length of the linker between the C-terminus of the protein of interest and MNase is 33 aa (GRRIPGLIKDYKDHDGDYKDHDIDYKDDDDKAA). ChEC-tagging vectors with the HIS3MX6 and TRP1 markers (pGZ109 and pGZ110), derived from pFA6a-3HA-HIS3MX6^55^ (Addgene #41600) and pFA6a-3HA-TRP1^55^ (Addgene #41595), were also created but not used in this study. These vectors retain compatibility with the F2/R1 primer pairs commonly used for epitope tagging of endogenous yeast genes. The ChEC-tagging short linker vector pGZ173 was constructed as above, except that the 3xFLAG tag was excluded from the inserted PCR amplicon. The linker length of this vector is 8 aa (GRRIPGLI). Yeast ChEC strains were created by transformation with ChEC cassettes amplified from pGZ108 using gene-specific F2/R1 primers (http://yeastgfp.yeastgenome.org/yeastGFPOligoSequence.txt). The *REB1* promoter-3FLAG-MNase-NLS construct (pGZ136) was created by gene synthesis (Operon) and cloned into the XhoI and EcoRI sites of pRS406 for integration at *URA3.* A codon-optimized SV40 NLS (PPKKKRKV) was added to the C-terminus of MNase by PCR prior to ligation into pRS406. The vector for expressing N-terminally MNase-tagged Reb1 (pGZ172) was constructed by Gibson assembly^56^ of PCR amplicons encoding the *REB1* promoter, MNase-3FLAG, and the *REB1* ORF into the SacI site of pRS413. Following transformation of yeast with pGZ172, the chromosomal copy of *REB1* was deleted using a kanMX6 deletion cassette amplified from pFA6a-kanMX6 with an F2/R1 primer pair. Plasmids and yeast strains used in this study are listed in Supplementary Tables 1 and 2, respectively.

### ChEC

For each ChEC experiment, a 50 mL culture was grown to an OD_600_ of 0.5-0.7 at 30°C in YPD. Cells were pelleted at 3,000 x *g* and washed three times with 1 mL Buffer A (15 mM Tris pH 7.5, 80 mM KCl, 0.1 mM EGTA, 0.2 mM spermine, 0.5 mM spermidine, 1X Roche cOmplete EDTA-free mini protease inhibitors, 1 mM PMSF), centrifuging as above between washes. Cells were resuspended in 600 μL Buffer A containing 0.1% digitonin and permeabilized at 30°C for 5 min. CaCl_2_ was added to 2 mM and ChEC digests were performed at 30°C. At each time point, a 100 μL aliquot of the digest was transferred to a tube containing 100 μL 2X stop buffer (400 mM NaCl, 20 mM EDTA, 4 mM EGTA) and 1% SDS. Protein was then digested with 80 μg proteinase K at 55°C for 30 m. Nucleic acids were extracted with an equal volume of phenol/chloroform/isoamyl alcohol and precipitated with 2.5 volumes 100% ethanol and 30 μg Glycoblue (Ambion). Pellets were washed once with 1 mL 100% ethanol, dried, and resuspended in 30 μL 0.1X TE buffer, pH 8.0. RNA was digested with 10 μg RNase A at 37°C for 20 m.

### Western blotting

Yeast cells were grown to an OD_600_ of 0.4-0.5 in YPD and whole cell extract (WCE) was prepared from 2 OD_600_ as described^57^. Antibodies used for western blotting were rabbit anti-ɣH2A (H2A phospho S129) (Abcam ab15083, 1:500) and rabbit anti-H2A (Millipore 07-146, 1:1000). Images were obtained using the Licor Odyssey and bands were quantified using Image Studio Lite (Licor). The ratio of ɣH2A/total H2A in the WT WCE was set to 1.0 and ɣH2A/total H2A ratios in the ChEC strains were expressed relative to this value.

### Library preparation and sequencing

Prior to generating sequencing libraries, ChEC DNA was subjected to size-selection using Ampure XP beads (Agencourt). A beads:sample ratio of 2.5:1 (v:v) was used and the unbound fraction, containing small DNA fragments, was extracted to remove RNase and precipitated as above, and used for library preparation. Sequencing libraries were constructed as described^46,58^, except that KAPA polymerase was used for library amplification. Libraries were sequenced for 25 cycles in paired-end mode on the Illumina HiSeq 2500 platform at the Fred Hutchinson Cancer Research Center Genomics Shared Resource. Paired-end fragments were mapped to the sacCer3/V64 genome build using Novoalign (Novocraft) as described to generate SAM files^46,59^. For visualization of ChEC-seq tracks, data were normalized as follows. The number of fragment ends corresponding to each base position in the genome was divided by the total number of read ends mapped. This accounts for differences in sequencing depth across samples. Read end-normalized counts/bp were then scaled by multiplication of each position with the total number of mapped bases in that sample. Normalization was performed using a custom perl script (pairs_single_end_sizes.pl in Supplementary File 1).

### Comparison of ChEC-seq data to ChIP data

The sum of ChEC cleavages within 100 bp windows centered on each peak midpoint was determined using the *bedmap* feature of the BEDOPS suite^60^. Abf1 and Reb1 ORGANIC peaks (10 m MNase, 80 mM salt) were from Kasinathan *et al*^12^, Rap1 ChIP-exo peaks were from Rhee and Pugh^8^, and Abf1, Rap1, and Reb1 X-ChIP-chip peaks were from MacIsaac *et al*^45^. An equal number of random sites for each peak set was generated using the *random* feature of the BEDTools suite, filtered to exclude any high-scoring or low-scoring sites^61^. Overlap of ChEC-seq and ChIP peaks was performed using BEDTools *intersect* using 100 bp windows centered around the motif match midpoint for ChEC-seq peaks or the peak midpoint for ChIP peaks. To analyze X-ChIP signal at Abf1 ChEC-seq sites, we performed Abf1 MNase-X-ChIP as described^13^. Average plots were generated using bedgraph files with a custom perl script with a shell wrapper (average_plot.pl and window.sh in Supplementary File 1).

### Peak calling

The maximum signal across all time points for each base position in the genome was determined using a custom perl script (combine_chec_bed.pl in Supplementary File 1) that output a single bedgraph file with the maximum value for each position. Peaks were called on this bedgraph file using a genome-wide thresholding approach using a custom perl script with a shell wrapper (threshold_bed.pl and call_peaks.sh in Supplementary File 1). To be included in a peak, a base position was required to have a value at least 10 times the genome-wide average. Positions exceeding the specified threshold within 30 bp of one another were merged into a single peak (an interpeak distance of 30). Thresholds used were: Abf1, 10.02165; Rap1, 9.24372; Reb1, 8.17298; free MNase, 10.57292. Reproducibility of peak occupancies was assessed by Spearman correlation. To calculate an FDR for each dataset, TF and free MNase peaks were overlapped using BEDOPS --*element-of* and the number of overlapping peaks was divided by the total number of TF peaks. FDRs calculated were: Abf1-MNase, 2.37%; Rap1-MNase, 3.71%; Reb1-MNase, 2.42%. Information on called peaks is given in Supplementary Data 1.

### Temporal analysis of ChEC-seq data

The sum of ChEC cleavages within 50 bp windows centered on each peak midpoint was determined. Rows were then Z-score transformed to allow comparison of ChEC-seq signal across rows (multiple time points). This analysis was performed with a custom perl script (chec_heatmap.pl in Supplementary File 1). Matrices were clustered with Gene Cluster 3.0^62^ and visualized with Java Treeview.

### Motif analysis

FASTA sequences were obtained for each 50 bp window surrounding each peak midpoint using BEDTools *fastaFromBed.* FASTA sequences were then scored using a custom perl script (pssm_scorer.pl in Supplementary File 1) implementing the FIMO algorithm^63^ and using previously determined PFMs downloaded from ScerTF^64^. FASTA sequences were uploaded to the MEME-ChIP^65^ web server for *de novo* motif discovery. Logos were generated with LogOddsLogo^66^, using the yeast GC content option.

### Cleavage pattern analysis

A custom perl script with a shell wrapper (average_plot_ends.pl and window_ends.sh in Supplementary Software) was used to determine average fragment end counts from pairs.bed files (created using pairs2bed.sh, Supplementary File 1) at each position in a 100 bp window surrounding each motif match center. Cleavage data from the time point with maximal signal for each factor and class of sites in the Figure 3 heatmaps were used for average analysis. Cleavage data from 30 s and 5 m time points were used for average analysis of SL and N-terminal cleavage. Counts were normalized by multiplication of each position by the size of the budding yeast genome (taken here to be 12,495,000 bp) divided by the number of fragments mapped for a given time point. Normalized free MNase counts were then subtracted from TF-MNase counts.

### DNA shape analysis

FASTA sequences generated from the previously generated 100 bp windows centered on motif match midpoints were used as input for our DNAshape method^67^ for high-throughput prediction of DNA structural features. A set of 100 bp windows generated by BEDTools *random* not overlapping high-scoring or low-scoring sites and equal in number to the low-scoring sites for each factor was used as the random control for DNA shape analysis. The resulting patterns for minor groove width (MGW), Roll, propeller twist (ProT), and helix twist (HelT) were analyzed using the framework of the motif database TFBSshape^68^. Statistical comparisons between DNA shape profiles were performed using a Pearson correlation coefficient (PCC) and Kolmogorov-Smirnov (KS) test with the null hypothesis that the distributions of DNA shape profiles derived from sequences containing high-scoring and low-scoring motifs are identical. PCCs close to 1 and large KS *p* values, therefore, indicate similar DNA shape profiles.

### Multiple linear regression

To test if DNA sequence or shape are more similar between sequences containing high-scoring and low-scoring motifs, we trained two models based on L2-regularized multiple linear regression, one model using DNA sequence and the other model using four DNA shape features: MGW, ProT, Roll, and HelT. The sequence was encoded in binary features, whereas the shape features were normalized between 0 and 1, as previously described^49^. Using the sequence-based and shape-based models, values for the area under the receiver operating curve (AUROC) were calculated for the classification between sequences containing high-scoring and low-scoring motifs and compared to the classification of sequences containing high-scoring motifs and random sequences. An AUROC value of 0.5 indicates a random classifier.

### Data availability

Sequencing data have been deposited in GEO under accession number GSE67453.

## Acknowledgements

We thank Jorja Henikoff for assistance with data analysis and Lin Yang for helpful suggestions and assistance with machine learning analysis. This work was supported by NIH grants T32CA009657 (G.E.Z.), F30CA186458 (S.K.), R01GM106056 (R.R.), R01ES020116, and the Howard Hughes Medical Institute (S.H.).

## Author contributions

G.E.Z. adapted ChEC to a sequencing readout and designed the study with S.H.; G.E.Z. performed ChEC-seq experiments; S.K. performed Abf1 MNase-X-ChIP and wrote software; G.E.Z. and S.K. analyzed ChEC-seq data; B.X. and R.R. performed DNA shape and machine learning analyses; G.E.Z. wrote the paper with input from all authors.

## References

1 Zentner, G. E. & Henikoff, S. High-resolution digital profiling of the epigenome. Nature Reviews Genetics 15, 814–827 (2014).

2 Jackson, V. Formaldehyde Cross-Linking for Studying Nucleosomal Dynamics. Methods 17, 125–139 (1999).

3 Poorey, K. et al. Measuring Chromatin Interaction Dynamics on the Second Time Scale at Single-Copy Genes. Science 342, 369–372 (2013).

4 Chen, J. et al. Single-Molecule Dynamics of Enhanceosome Assembly in Embryonic Stem Cells. Cell 156, 1274–1285 (2014).

5 Fan, X. & Struhl, K. Where Does Mediator Bind In Vivo? PLoS ONE 4, e5029 (2009).

6 Teytelman, L., Thurtle, D. M., Rine, J. & van Oudenaarden, A. Highly expressed loci are vulnerable to misleading ChIP localization of multiple unrelated proteins. Proc. Natl. Acad. Sci. USA 110, 18602–18607 (2013).

7 Park, D., Lee, Y., Bhupindersingh, G. & Iyer, V. R. Widespread Misinterpretable ChIP-seq Bias in Yeast. PLoS ONE 8, e83506 (2013).

8 Rhee, H. S. & Pugh, B. F. Comprehensive Genome-wide Protein-DNA Interactions Detected at Single-Nucleotide Resolution. Cell 147, 1408–1419 (2011).

9 Skene, P. J., Hernandez, A. E., Groudine, M. & Henikoff, S. The nucleosomal barrier to promoter escape by RNA polymerase II is overcome by the chromatin remodeler Chd1. eLife 3, e02042 (2014).

10 He, Q., Johnston, J. & Zeitlinger, J. ChIP-nexus enables improved detection of in vivo transcription factor binding footprints. Nat Biotech 33, 395–401 (2015).

11 Teytelman, L. et al. Impact of Chromatin Structures on DNA Processing for Genomic Analyses. PLoS ONE 4, e6700 (2009).

12 Kasinathan, S., Orsi, G. A., Zentner, G. E., Ahmad, K. & Henikoff, S. High-resolution mapping of transcription factor binding sites on native chromatin. Nat. Methods 11, 203–209 (2014).

13 Zentner, G. E., Tsukiyama, T. & Henikoff, S. ISWI and CHD chromatin remodelers bind promoters but act in gene bodies. PLoS Genet. 9, e1003317 (2013).

14 van Steensel, B. & Henikoff, S. Identification of in vivo DNA targets of chromatin proteins using tethered Dam methyltransferase. Nat. Biotechnol. 18, 424–428 (2000).

15 Vogel, M. J., Peric-Hupkes, D. & van Steensel, B. Detection of in vivo protein-DNA interactions using DamID in mammalian cells. Nat. Protocols 2, 1467–1478 (2007).

16 Filion, G. J. et al. Systematic Protein Location Mapping Reveals Five Principal Chromatin Types in Drosophila Cells. Cell 143, 212–224 (2010).

17 Germann, S., Juul-Jensen, T., Letarnec, B. & Gaudin, V. DamID, a new tool for studying plant chromatin profiling in vivo, and its use to identify putative LHP1 target loci. Plant J. 48, 153–163 (2006).

18 Steglich, B., Filion, G., van Steensel, B. & Ekwall, K. The inner nuclear membrane proteins Man1 and Ima1 link to two different types of chromatin at the nuclear periphery in S. pombe. Nucleus 3, 77–87 (2012).

19 Schuster, E. et al. DamID in C. elegans reveals longevity-associated targets of DAF-16/FoxO. Mol. Syst. Biol. 6, 399 (2010).

20 Wang, H., Mayhew, D., Chen, X., Johnston, M. & Mitra, R. D. “Calling Cards” for DNA-Binding Proteins in Mammalian Cells. Genetics 190, 941–949 (2012).

21 Schmid, M., Durussel, T. & Laemmli, U. K. ChIC and ChEC: Genomic Mapping of Chromatin Proteins. Mol. Cell 16, 147–157 (2004).

22 Merz, K. et al. Actively transcribed rRNA genes in S. cerevisiae are organized in a specialized chromatin associated with the high-mobility group protein Hmo1 and are largely devoid of histone molecules. Genes Dev. 22, 1190–1204 (2008).

23 Schmid, M. et al. Nup-PI: The Nucleopore-Promoter Interaction of Genes in Yeast. Mol. Cell 21, 379–391 (2006).

24 Dunn, T., Gable, K. & Beeler, T. Regulation of cellular Ca2+ by yeast vacuoles. J. Biol. Chem. 269, 7273–7278 (1994).

25 Lyng, F. M., Jones, G. R. & Rommerts, F. F. G. Rapid Androgen Actions on Calcium Signaling in Rat Sertoli Cells and Two Human Prostatic Cell Lines: Similar Biphasic Responses Between 1 Picomolar and 100 Nanomolar Concentrations. Biol. Reprod. 63, 736–747 (2000).

26 Mountjoy, K. G., Kong, P. L., Taylor, J. A., Willard, D. H. & Wilkison, W. O. Melanocortin receptor-mediated mobilization of intracellular free calcium in HEK293 cells. Physiol. Genomics 5, 11–19 (2001).

27 Grienberger, C. & Konnerth, A. Imaging Calcium in Neurons. Neuron 73, 862–885 (2012).

28 Schmidt, T., Schirra, C., Matti, U., Stevens, D. R. & Rettig, J. Snapin accelerates exocytosis at low intracellular calcium concentration in mouse chromaffin cells. Cell Calcium 54, 105–110 (2013).

29 Seurin, D., Lombet, A., Babajko, S., Godeau, F. & Ricort, J.-M. Insulin-Like Growth Factor Binding Proteins Increase Intracellular Calcium Levels in Two Different Cell Lines. PLoS ONE 8, e59323 (2013).

30 Rhode, P. R., Elsasser, S. & Campbell, J. L. Role of multifunctional autonomously replicating sequence binding factor 1 in the initiation of DNA replication and transcriptional control in Saccharomyces cerevisiae. Mol. Cell. Biol. 12, 1064–1077 (1992).

31 Reed, S. H., Akiyama, M., Stillman, B. & Friedberg, E. C. Yeast autonomously replicating sequence binding factor is involved in nucleotide excision repair. Genes Dev. 13, 3052–3058 (1999).

32 Moehle, C. M. & Hinnebusch, A. G. Association of RAP1 binding sites with stringent control of ribosomal protein gene transcription in Saccharomyces cerevisiae. Mol. Cell. Biol. 11, 2723–2735 (1991).

33 Lustig, A., Kurtz, S. & Shore, D. Involvement of the silencer and UAS binding protein RAP1 in regulation of telomere length. Science 250, 549–553 (1990).

34 Lang, W. H. & Reeder, R. H. The REB1 site is an essential component of a terminator for RNA polymerase I in Saccharomyces cerevisiae. Mol. Cell. Biol. 13, 649–658 (1993).

35 Scott, E. W. & Baker, H. V. Concerted action of the transcriptional activators REB1, RAP1, and GCR1 in the high-level expression of the glycolytic gene TPI. Mol. Cell. Biol. 13, 543–550 (1993).

36 Graham, I. R. & Chambers, A. A Reb1p-binding site is required for efficient activation of the yeast RAP1 gene, but multiple binding sites for Rap1p are not essential. Mol. Microbiol. 12, 931–940 (1994).

37 Goetze, H. et al. Alternative Chromatin Structures of the 35S rRNA Genes in Saccharomyces cerevisiae Provide a Molecular Basis for the Selective Recruitment of RNA Polymerases I and II. Mol. Cell. Biol. 30, 2028–2045 (2010).

38 Reiter, A. et al. The Reb1 homologue Ydr026c/Nsi1 is required for efficient RNA polymerase I termination in yeast. 31, 3480–3493 (2012).

39 Hartley, P. D. & Madhani, H. D. Mechanisms that Specify Promoter Nucleosome Location and Identity. Cell 137, 445–458 (2009).

40 Ganapathi, M. et al. Extensive role of the general regulatory factors, Abf1 and Rap1, in determining genome-wide chromatin structure in budding yeast. Nucleic Acids Res. 39, 2032–2044 (2011).

41 van Bakel, H. et al. A Compendium of Nucleosome and Transcript Profiles Reveals Determinants of Chromatin Architecture and Transcription. PLoS Genet. 9, e1003479 (2013).

42 Biggin, Mark D. Animal Transcription Networks as Highly Connected, Quantitative Continua. Dev. Cell 21, 611–626 (2011).

43 Fisher, W. W. et al. DNA regions bound at low occupancy by transcription factors do not drive patterned reporter gene expression in Drosophila. Proceedings of the National Academy of Sciences of the United States of America 109, 21330–21335 (2012).

44 Ghaemmaghami, S. et al. Global analysis of protein expression in yeast. Nature 425, 737–741 (2003).

45 MacIsaac, K. et al. An improved map of conserved regulatory sites for Saccharomyces cerevisiae. BMC Bioinformatics 7, 113 (2006).

46 Henikoff, J. G., Belsky, J. A., Krassovsky, K., MacAlpine, D. M. & Henikoff, S. Epigenome characterization at single base-pair resolution. Proceedings of the National Academy of Sciences of the United States of America 108, 18318–18323 (2011).

47 Matot, B. et al. The orientation of the C-terminal domain of the Saccharomyces cerevisiae Rap1 protein is determined by its binding to DNA. Nucleic Acids Res. 40, 3197–3207 (2012).

48 Feeser, E. A. & Wolberger, C. Structural and Functional Studies of the Rap1 C-Terminus Reveal Novel Separation-of-Function Mutants. J. Mol. Biol. 380, 520–531 (2008).

49 Abe, N. et al. Deconvolving the Recognition of DNA Shape from Sequence. Cell 161, 307–318 (2015).

50 Zhou, T. et al. Quantitative modeling of transcription factor binding specificities using DNA shape. Proc. Natl. Acad. Sci. USA 112, 4654–4659 (2015).

51 Slutsky, M. & Mirny, L. A. Kinetics of Protein-DNA Interaction: Facilitated Target Location in Sequence-Dependent Potential. Biophys. J. 87, 4021–4035 (2004).

52 Neph, S. et al. An expansive human regulatory lexicon encoded in transcription factor footprints. Nature 489, 83–90 (2012).

53 Sung, M.-H., Guertin, Michael J., Baek, S. & Hager, Gordon L. DNase Footprint Signatures Are Dictated by Factor Dynamics and DNA Sequence. Mol. Cell 56, 275–285 (2014).

54 Sander, J. D. & Joung, J. K. CRISPR-Cas systems for editing, regulating and targeting genomes. Nat Biotech 32, 347–355 (2014).

55 Longtine, M. S. et al. Additional modules for versatile and economical PCR-based gene deletion and modification in Saccharomyces cerevisiae. Yeast 14, 953–961 (1998).

56 Gibson, D. G. et al. Enzymatic assembly of DNA molecules up to several hundred kilobases. Nat. Methods 6, 343–345 (2009).

57 Kushnirov, V. V. Rapid and reliable protein extraction from yeast. Yeast 16, 857–860 (2000).

58 Zentner, G. E. & Henikoff, S. Mot1 redistributes TBP from TATA-containing to TATA-less promoters. Mol. Cell. Biol. 33, 4996–5004 (2013).

59 Krassovsky, K., Henikoff, J. G. & Henikoff, S. Tripartite organization of centromeric chromatin in budding yeast. Proceedings of the National Academy of Sciences of the United States of America 109, 243–248 (2012).

60 Neph, S. et al. BEDOPS: high-performance genomic feature operations. Bioinformatics 28, 1919–1920 (2012).

61 Quinlan, A. R. & Hall, I. M. BEDTools: a flexible suite of utilities for comparing genomic features. Bioinformatics 26, 841–842 (2010).

62 de Hoon, M. J. L., Imoto, S., Nolan, J. & Miyano, S. Open source clustering software. Bioinformatics 20, 1453–1454 (2004).

63 Grant, C. E., Bailey, T. L. & Noble, W. S. FIMO: scanning for occurrences of a given motif. Bioinformatics 27, 1017–1018 (2011).

64 Spivak, A. T. & Stormo, G. D. ScerTF: a comprehensive database of benchmarked position weight matrices for Saccharomyces species. Nucleic Acids Res. 40, D162-D168 (2012).

65 Machanick, P. & Bailey, T. L. MEME-ChIP: motif analysis of large DNA datasets. Bioinformatics 27, 1696–1697 (2011).

66 Yu, Y.-K., Capra, J. A., Stojmirović, A., Landsman, D. & Altschul, S. F. Log-odds sequence logos. Bioinformatics 31, 324–331 (2015).

67 Zhou, T. et al. DNAshape: a method for the high-throughput prediction of DNA structural features on a genomic scale. Nucleic Acids Res. 41, W56–W62 (2013).

68 Yang, L. et al. TFBSshape: a motif database for DNA shape features of transcription factor binding sites. Nucleic Acids Res. 42, D148–D155 (2014).

